# Artificial Light at Night Disrupts Circadian and Metabolic Gene Expression in the Green Anole Lizard (*Anolis carolinensis*): A Transcriptomic Analysis

**DOI:** 10.1101/2025.03.20.644270

**Authors:** Violeta Trejo-Reveles, Michele A. Johnson, Florissa F. Patterson, Grace E. Anderson, Zhou Wu, Alex R. Johnston, Simone L. Meddle

## Abstract

Artificial light at night (ALAN) disrupts natural light-dark cycles, posing ecological challenges for wildlife in urban areas. Here we investigated the effects of ALAN on gene expression in the brain, liver, and skin of green anole lizards (*Anolis carolinensis*) whose urban populations are increasingly exposed to light pollution. To identify genetic pathways impacted by ALAN exposure we analysed expression of genes associated with circadian and metabolic regulation at midday, midnight and at midnight with artificial light. Differential expression analysis revealed that clock-related genes (*PER1*, *NR1D1*, *CRY2*) were significantly altered in the brain, liver, and skin following ALAN treatment and genes involved in glucagon regulation (*GCG*) and lipid metabolism (*NOCT*) were differentially expressed in the liver, indicating metabolic disruptions. Skin exhibited unique responses to ALAN suggesting that repair responses may be altered as genes related to cellular processes, such as wound healing, were upregulated under normal light and dark conditions. Our findings also show that ALAN disrupts core circadian genes, impacting physiological processes including hormone regulation, glucose homeostasis, and potentially reproductive cycles. This study provides the first transcriptomic evidence of the effects of light pollution on green anoles, highlighting the need to preserve natural light cycles in urban habitats. An interactive online database developed for this study allows further exploration of gene expression changes, to promote research on artificial light-polluted environments.

## Introduction

Light pollution, defined as the alteration of natural light-dark cycles by artificial light sources, has become a persistent environmental concern worldwide, particularly in urban areas. The disruption of a natural light-dark cycle can have detrimental effects on wildlife physiology and behaviour. These disruptions are best-studied in mammals. For example, it has been shown that foraging behaviour in the nocturnal Mongolian five-toed jerboa (*Allactaga sibirica*) is altered following exposure to artificial light at night (ALAN; Zhang et al., 2020) and other nocturnal rodents such as wild mice (*Mus musculus*) decreased their normal activity levels when artificial light was present (Oosthuizen et al., 2024). ALAN has also been reported to have a negative effect on homeostasis in the spiny mouse (*Acomys cahirinus*) as chronic elevated cortisol levels and higher mortality rates were observed (Vardi-Naim et al., 2022), and in laboratory mice ALAN prevented weight gain (Melendez-Fernandez et al., 2023). Population size in wild mammalian species can be heavily impacted by urbanisation and light pollution through effects on the reproductive clock. For example, in tammar wallabies (*Macropus eugenii),* ALAN suppressed the melatonin levels and delayed births (Robert et al., 2015).

ALAN has also been reported to affect other groups of wild vertebrates. Studies have shown that artificial light not only affects bird behaviour, but also health and reproduction by altering physiology and activity cycles (Amichai & Kronfeld-Schor 2019; Dominoni et al., 2013). Furthermore, illumination at night can severely impact bird migration patterns, as well as avian perceptions of habitat quality as illuminated areas are avoided (Adams et al., 2021). Studies in amphibians have shown that artificial light exposure can change American toad (*Anaxyrus americanus*) activity cycles (Dananay & Bernard, 2018) with the potential to influence the ecosystem equilibrium as different phenotypes can alter predator perception and amphibian population sizes (Shidemantle et al., 2022). ALAN has also been shown to affect offspring behaviour in fish. Zebrafish (*Danio rerio*) that were exposed to constant artificial light, not only showed altered behaviour, but F1 offspring born from ALAN-exposed mothers displayed less frequent movement and shorter movement distances despite never being exposed to ALAN themselves (Lim et al., 2024).

Reptiles are frequently exposed to light pollution in urban habitats, and the effects of artificial light on their physiology and behaviour is less understood. The green anole lizard (*Anolis carolinensis*) is a valuable species for studying the effects of ALAN as their behaviour and physiological processes, such as circadian rhythms, thermoregulation, and reproduction, are strongly influenced by light, including ALAN (Bayard 1974; Taylor et al., 2022). Green anoles are small, diurnal, arboreal lizards that are commonly found in habitats ranging from dense forests to urban areas and are native to the south eastern United States. Effects of ALAN in green anoles include increased nocturnal foraging and display behaviour, reduced daytime activity, and changes in reproductive organ size (Taylor et al., 2022) and such behavioural shifts are likely mediated by changes at the genetic level. Biological internal clocks regulate daily cycles of physiological activity, and are controlled by complex genetic networks that respond to external light cues. Disruption of these natural light-dark cycles by ALAN can interfere with the circadian system through the expression of circadian rhythm-related genes to impact a range of biological processes (Ouyang et al., 2018; Taylor et al., 2022; Thawley & Kolbe, 2020). Previous notable work in reptiles has reported the effect of circadian rhythm disruption on metabolism and energy regulation as ALAN has been shown to impact liver clock gene expression e.g., *PER1* and *GCG*, leading to metabolic imbalances, weight gain and glucose intolerance (Guan & Lazar 2022; Park et al., 2019).

Valuable insights into the underlying genetic and physiological impacts of altered light-dark cycles by artificial light exposure on the disruption of core clock genes, including *PER1*, *CRY1*, *NR1D1*, and *BMAL1*, has been extensively studied in laboratory mice (Bugge et al., 2012; Sato et al., 2004). ALAN exposure in mice has been reported to affect the rhythmic gene expression of *PER1* and *NR1D1*, which play key roles in maintaining circadian stability. These genes act as transcriptional regulators that link light exposure to physiological rhythms (Chauvet et al., 2016). Disruption by ALAN can cause “phase shifts” in feeding, energy metabolism, and sleep-wake cycles, leading to desynchronization between internal rhythms and the external environment. For example, nocturnal exposure to light disrupts *CRY2* and *PER2* expression which results in altered sleep patterns, hormonal imbalances, and alterations in glucose metabolism (Kalsbeek et al., 2010; Grunst et al., 2023). Other circadian-regulated genes, such as *NOCT*, are involved in lipid metabolism and *NOCT* expression is altered in response to ALAN, which results in changes in lipid storage and transport (Kulshrestha et al., 2023).

Recent advances in transcriptomic analysis allow for a more detailed investigation into the molecular effects of artificial light exposure in green anole lizards. Using the latest annotated green anole genome (AnoCar2.0v2), gene expression profiling was used in this current study to identify differentially expressed genes (DEGs) associated with artificial light exposure, highlighting potential molecular mechanisms by which ALAN affects behaviour and physiology. Green anoles also exhibit unique adaptations that make them suitable for studying photoreception beyond ocular tissues. In addition to retinal opsins, anoles express photoreceptive proteins in extra-retinal tissues, such as the skin and brain, potentially allowing them to detect environmental light changes directly through these structures (Porter et al., 2011; Perez et al., 2019). These expression patterns suggest that there is a complex sensory network that could contribute to light-dependent behaviour and physiology in urban environments.

Given the ecological importance of lizards and the potential implications of light pollution for health and behaviour, this study investigated the effects of ALAN on gene expression in the green anole brain, liver, and skin. By analysing changes in gene expression across tissues, specific genetic pathways regulating circadian rhythms and metabolic processes were found to be affected by ALAN. Understanding these molecular responses provides a foundation for assessing the broader ecological impacts of light pollution on vertebrates, as well as informing conservation strategies to mitigate the effects of urbanisation on wildlife.

## Materials and Methods

### Animal capture and housing

Twenty-four free-living adult green anole lizards (*Anolis carolinensis*; twelve of each sexwere captured in the breeding season, in May 2024, on the urban campus of Trinity University, San Antonio, Texas, USA, during daylight hours. Green anole lizards were collected by using a dental floss loop attached to an extendable fishing pole or by hand and were transported individually to the Trinity University vivarium in cotton bags. On the day of capture, the body mass of each anole was measured to the nearest 0.1 g using a Pesola spring scale and snout-vent length measured to the nearest mm using a clear plastic ruler (Males: range 52-69mm, average 62mm; Females: range 51-57mm, average 55mm). Each individual was given a unique identification number on the lower jaw using a non-toxic permanent marker. Each anole was then randomly assigned to one of three treatment groups: Midday, Midnight, or ALAN. Sex was determined by the presence of a dewlap; four anoles of each sex were assigned to each treatment group.

All green anoles were housed in the Trinity University vivarium following standard anole care procedures for a minimum of four days prior to tissue collection (Sanger et al., 2008). Pairs of anoles (one of each sex) were assigned to the same treatment and housed together in large Kritter Keeper™ cages (37.5 x 21.0 x 28.0 cm^3^; Lee’s Aquarium and Pet Products, San Marcos, CA, USA). Cages contained Zilla Green Terrarium Liner™ (Zilla, Franklin, WI, USA), 2 PVC pipe perches, a wire mesh hammock, and a nest box where females could lay eggs (i.e., a plastic flower pot with moist sphagnum moss). Cages were misted with water daily to provide drinking water, and each anole was fed approximately every other day between 12:00 and 18:00. At each feeding, two or three crickets were dusted with Zoo Med Repti Calcium™ supplement (Zoo Med Laboratories, Inc., San Luis Obispo, CA, USA).

In the vivarium, green anoles were housed together in one room for the Midday and Midnight treatments, and housed in an adjacent room for the ALAN treatment. The humidity and temperature ranged from 51 – 62 %, and 25.1 - 28.1 °C respectively, with similar conditions in both rooms. All cages were kept under standard lighting conditions on a 12.5 light: 11.5 dark cycle (room ceiling lights on at 06:00) to mimic the natural light-dark cycle for the month of May in San Antonio, Texas, USA. Two Reptisun 5.0 UVB light bulbs (Zoo Med Laboratories, Inc., San Luis Obispo, CA, USA; emission peaks at 410, 440, 550 and 580 nm and broadband emission centred at 350 nm) were positioned over each cage to simulate the full spectrum of natural sunlight. To mimic daily dawn and dusk, room ceiling lights (32-watt GE T8 Starcoat ECO bulbs, GE, Boston, MA, USA; emission peak 450 nm and broadband emission centred at 600 nm) were switched on 30 min. before and 30 min. after the cage lights. Ceiling lights were switched off at 19:30 in both rooms, but in the ALAN treatment room, a street lamp (D802-LED 12 ʹʹ low-profile area light; Deco Lighting, Inc. Commerce, CA, USA) identical to those used for nocturnal lighting on Trinity University’s campus, was switched on. The street lamp was covered with black mesh deer cloth to provide a light intensity of 1.21 µmol / m / s (approx. 89.6 lux; SD = 0.14 µmol / m^2^ / s;) at a distance of 180 cm from the lizard cages in the ALAN room (see Taylor et al., 2022). This light intensity mimics the light intensity of nocturnal lighting on campus (1.33 µmol/m2; SD = 0.16 µmol /m^2^ / s (approx. 98.5 lux / s; Taylor et al., 2022). The streetlamp was switched off at 06:00, when the ceiling lights of the room were switched on. ALAN green anoles were maintained in this street light treatment for 3-5 days prior to cull and tissue collection.

### Tissue collection

Green anoles were rapidly decapitated without prior anaesthesia to avoid any confounding anaesthetic effects on RNA expression. Brain (containing the pineal gland), eyes, dorsal skin, ventral skin, liver, and testes or ovaries were collected in under 7 min. 33 s and flash frozen on dry ice and stored at −80 °C until they were shipped to BGI Genomics (San Jose, CA, USA). Midday treatment dissections were performed between 12:48 to 13:45, and Midnight and ALAN treatment from 22:30 to 00:42. Decapitation for the Midnight treatment group was performed in the dark under red torch light (HQRP, Harrison, NJ; emission peak in the red spectrum at 650 nm). To control for this additional illumination, ALAN treatment lizards were also illuminated by the same red torch light during decapitation.

### Gene expression analysis

Tissues were shipped on dry ice to BGI Genomics (San Jose, CA, USA) where they were processed for RNA extraction, library preparation and sequencing. The quality of the RNA was checked before proceeding to library preparation to ensure a RIN of at least 7.0. Samples that did not pass the filter were discarded from the analysis. Raw fastq files were processed by trimming using trimmomatic (http://www.usadellab.org/cms/?page=trimmomatic) with the following parameters: ILLUMINACLIP:TruSeq3-PE-2.fa:2:30:10; SLIDINGWINDOW:10:30; LEADING:28; TRAILING:28; MINLEN:75. After trimming the quality of the reads was checked using FASTQC (https://github.com/s-andrews/FastQC). Once the parameters were checked, reads were aligned using STAR aligner version 2.7.11a (https://github.com/alexdobin/STAR), with the ENSEMBL green anole (*Anolis carolinensis*) genome and gtf (AnoCar2.0v2, Anolis_carolinensis.AnoCar2.0v2.112.gtf) as references. BAM files were counted using featurecounts (https://subread.sourceforge.net/).

Downstream statistical analysis was performed on R version 4.4.0. using the package Deseq2 (<u>Love, et. al., 2014</u>). Volcano, PCA, heatmaps and boxplots were drawn using ggplot2 (https://ggplot2.tidyverse.org/), and GO enrichment analysis was performed using the package genekitr (Liu, et.al.,2023) Venn diagrams were generated using the library VennDiagram (https://r-graph-gallery.com/14-venn-diagramm). GO classification analysis was performed in Panther (https://www.pantherdb.org/), by feeding the ENSEMBL IDs and selecting the green anole genome (*Anolis carolinensis*) as a reference. The web App was developed using R shiny (https://shiny.posit.co/), using Plotly to make any interactive plot interactive (https://plotly.com/r/). Genes were considered differentially expressed if they showed a log fold change of more than or equal to 1 or less than or equal to −1 and a p adjusted value of less or equal to 0.05.

### Data availability

**Raw data can be found in: GEOXXXX. And any** datasets used during the current study are available from the corresponding author upon request.

## Results

This study provides, to our knowledge, the first transcriptomic analysis of a reptile species in response to ALAN exposure. As an initial approach, a PCA analysis was conducted to examine variability among groups by comparing samples from lizards collected at Midday, Midnight (no light), and exposed to ALAN (midnight with artificial light). From this analysis, three distinct clusters emerged: one comprising of liver, a second and larger cluster containing skin (both dorsal and ventral), testes, and ovary samples with a subtle separation between skin and gonad groups, and a third cluster formed by brain and eye samples (Figure 1). A cluster composed of the skin, testes, and ovary samples showed minimal differences between light conditions. ALAN and Midnight liver samples clustered closely, while samples collected during Midday appeared more dispersed. For the brain, a clustering pattern emerged between Midday and Midnight samples, regardless of light exposure, although some outliers could be identified. No clustering was observed for liver or skin within light treatments (see Appendix 1).

**Figure 1.**
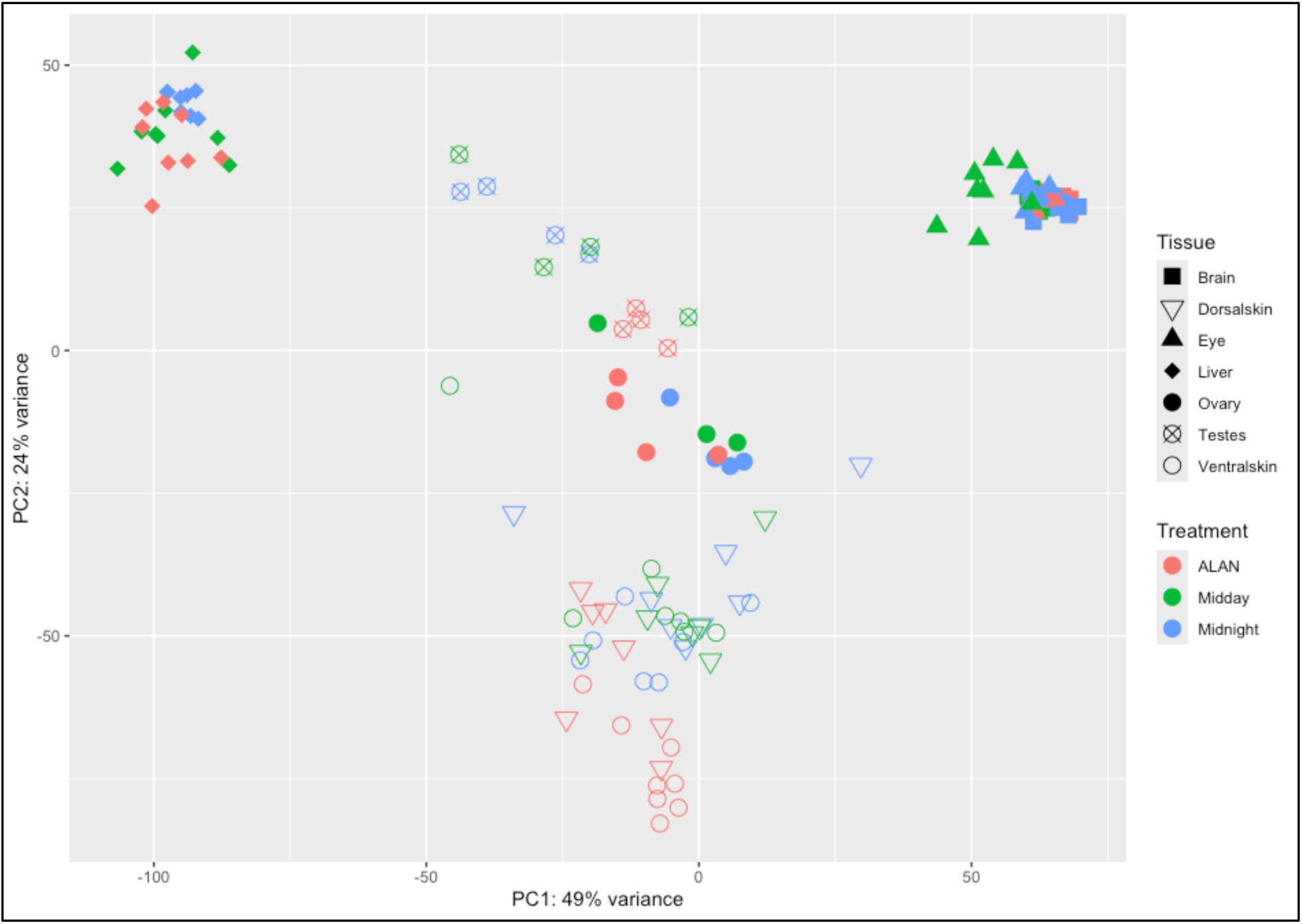
PCA plot comprising all samples from different tissues along with Midday, Midnight and ALAN treatments in green anole lizards. Shape depicts tissue, whilst colour illustrates treatment.

To further explore the effects of ALAN on different green anole tissues, ALAN samples were compared to those taken during the Midday and those taken at Midnight, to identify genes differentially expressed under “normal” day-night cycles and those differentially expressed under ALAN. Analysis was specifically focused on brain, which regulates circadian rhythms (Benca et al., 2009); liver, which reflects metabolic responses to artificial light (Thawley et al., 2020) and skin, which our unpublished data suggests may exhibit high photoreceptor variability (Trejo-Reveles et al., submitted). By comparing Midnight samples to those exposed to ALAN, circadian rhythm-related genes were found to be differentially expressed across tissues. Specifically, in the brain, the clock gene *PER1* (up-regulated;) and clock protein regulator *NR1D1* (down-regulated) emerged as the top differentially expressed genes. In the liver, genes associated with glucagon synthesis, such as *GCG* (up-regulated) and *GOT1L1* (down-regulated), were differentially expressed along with *PER1* (up-regulated) . In addition, circadian-related genes, including *DBP* (down-regulated), and *NR1D1* (down-regulated), were also differentially expressed in the skin. In the skin, a similar pattern to that observed in the brain and liver was identified, but additional genes, such as *CEBPD* and *M13A*, were also differentially expressed (Figure 2). When these results were compared to those obtained when samples were subjected to a natural light-dark exposure, *PER1* remained differentially expressed, particularly in the liver. Other genes, such as *NOCT*, *KLF9*, and *PPR31B*, also showed differential expression in this natural light comparison. Similar results were observed for the brain, where the majority of the genes that were DE in the Midday vs ALAN comparison were also DE in the Midday vs Midnight comparison. In the skin, a significantly lower number of genes were found DE when the tissue was not exposed to ALAN (Figure 2).

**Figure 2.**
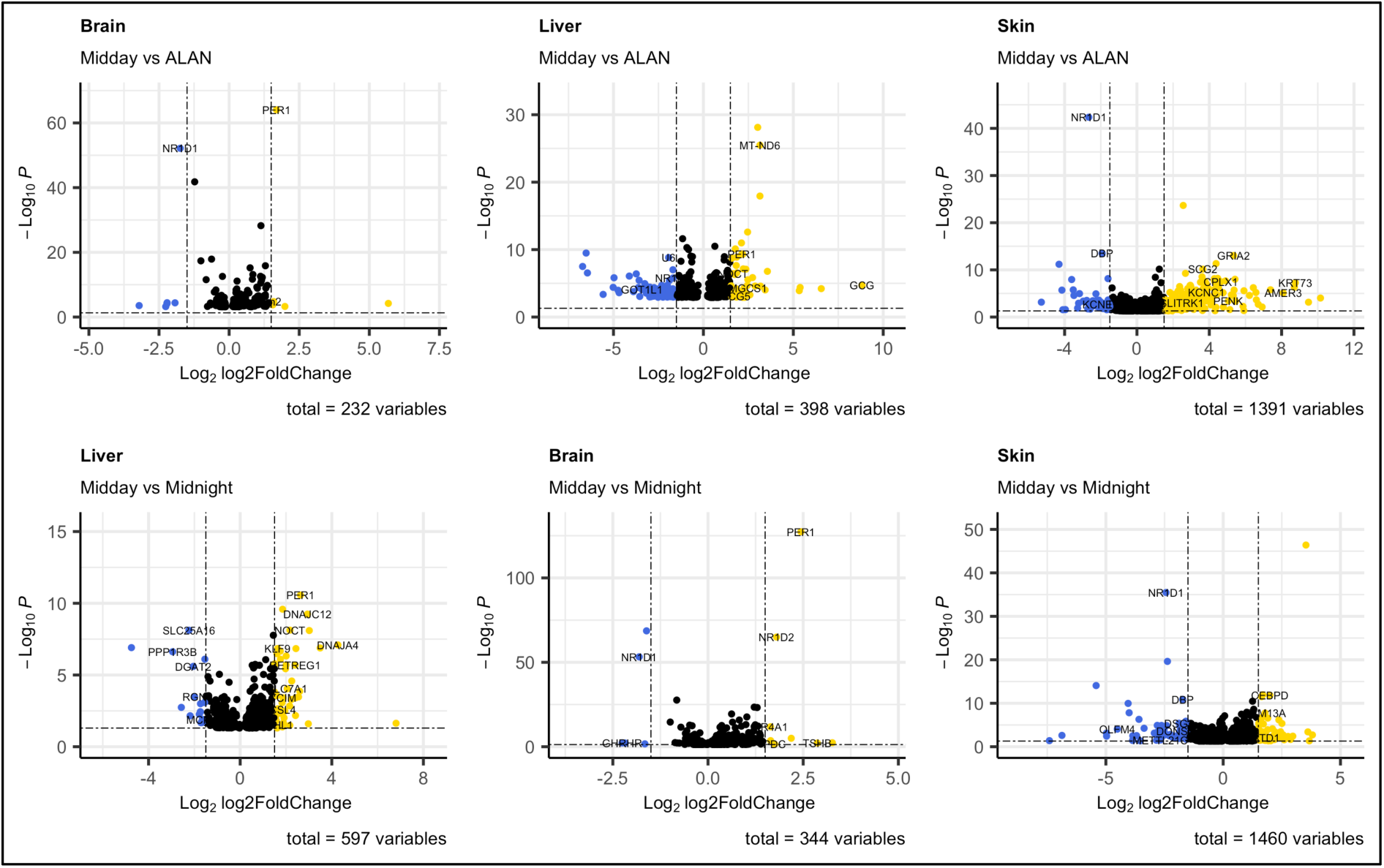
Volcano plots of differentially expressed genes (DEG). **A - C**: genes differentially expressed when comparing Midday vs ALAN; **D - F**: genes differentially expressed when comparing Midday vs Midnight. Comparisons are shown in the following order: Brain, Liver and Skin. Yellow dots depict up-regulated genes whilst blue dots represent down-regulated genes; black dots are non-significant genes. Significant genes were chosen based on Log fold change (</= −1 or > = 1) and p adjusted value (</= 0.05).

### Differences between ALAN exposure and natural light schedule

To further explore genes differentially expressed under a normal light-dark cycle compared to those under ALAN exposure, a Venn diagram analysis followed by gene ontology (GO) classification was conducted. The results obtained in the pairwise comparisons between Midday and Midnight and Midday and ALAN were compared and focused on both the shared DEGs and the DEGs that were exclusive to ALAN exposure. In the brain, only 26% of the DE genes were exclusive to the Midday vs. ALAN comparison (Figure 3). GO classification indicated that most of these genes were related to cellular process categories, including cell division and adhesion (see Appendix 2). Notably, *CRY2*, a circadian rhythm modulator was exclusively differentially expressed under ALAN exposure.

**Figure 3.**
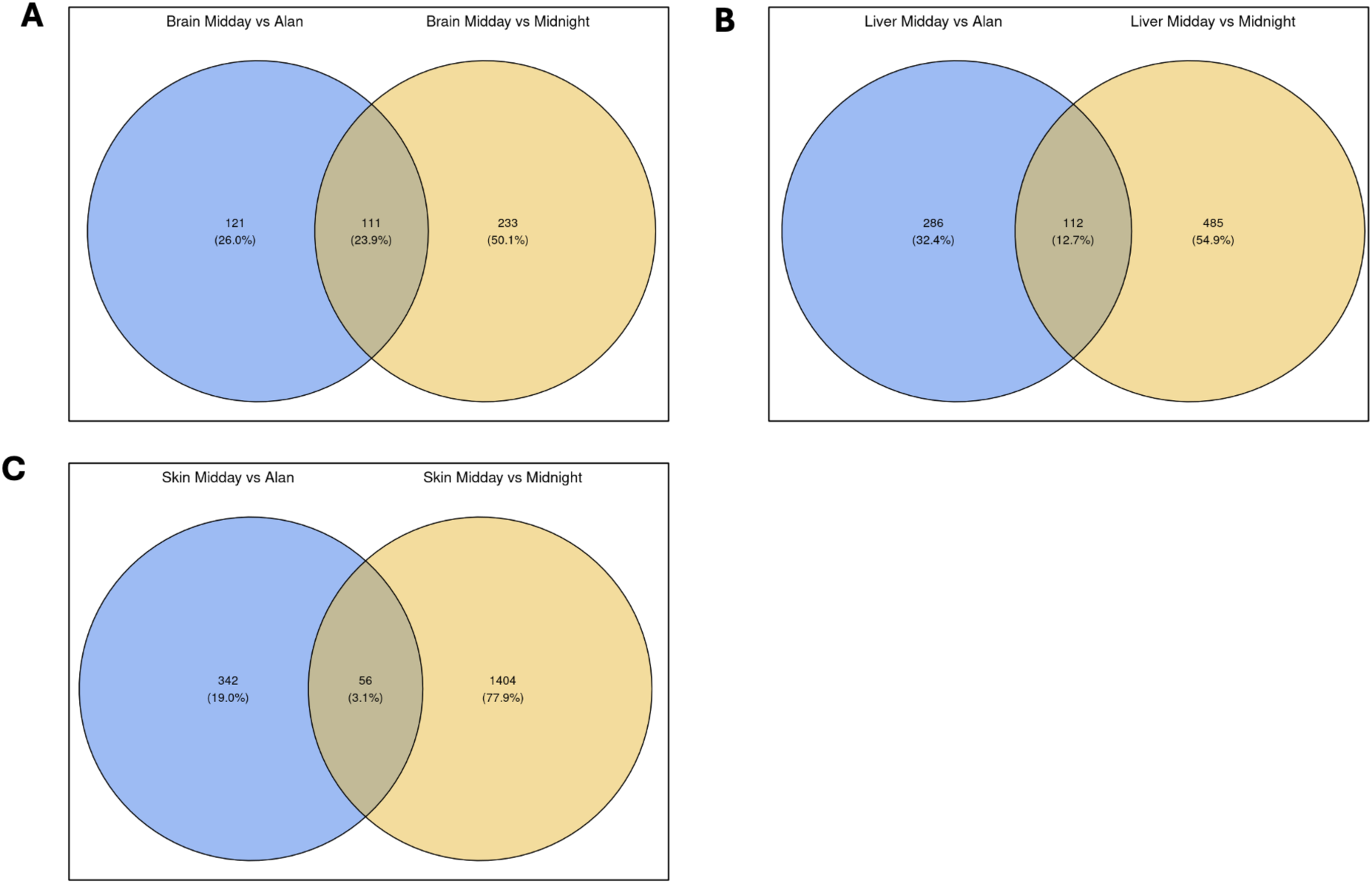
Venn diagrams of DEG for Midday vs ALAN or Midday vs Midnight comparisons. Each of the Venn diagrams show the similarity between comparisons for **A**: brain **B**: liver and **C**: skin.

**Figure 4.**
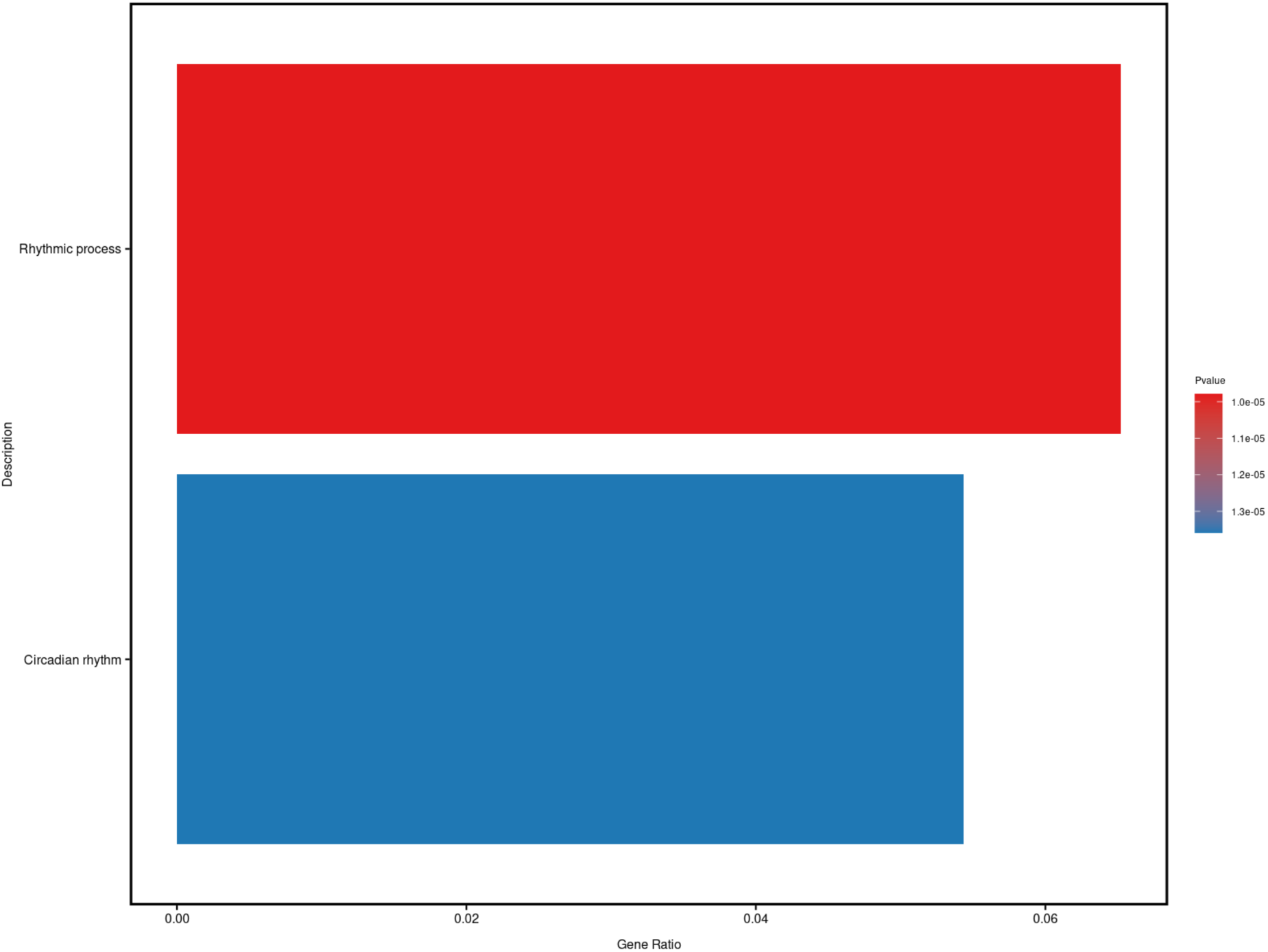
Enriched GO terms when comparing brain issue Midday vs ALAN. The colour is based on the p adjustment of the enrichment. Only two categories showed significant enrichment: rhythmic process and circadian rhythm.

In the liver, only 12.8% of DEG’s were shared between light-exposed and dark samples (Figure 3). Similar to the brain, shared terms in the liver were mostly related to cellular process, such as cell death and division. Some circadian-clock genes, including *PER1*, *CRY1*, and *PER2*, were commonly expressed. Genes exclusively DE in the liver under ALAN were primarily involved in metabolic processes, such as glycolysis. Interestingly, the F-box domain-containing protein, associated with circadian rhythm regulation, was DE in the Midday vs. ALAN comparison. *NOCT* (aka *Ccrn4l*), which is involved in the liver circadian clock and lipid metabolism (Kulshrestha et al., 2023), was downregulated under natural light-dark cycles but absent in DEG lists obtained under ALAN (Figure 3).

The largest discrepancy in DE genes was observed in the skin when comparing ALAN exposure to the other treatment groups. Only 3% of DEG’s were shared across comparisons, while a striking 78% were exclusive to ALAN exposure. Similar to the results observed for brain and liver, circadian clock-related genes, such as *CRY1*, *CRY2*, and *PER1*, were among the 56 shared genes. In the DE list exclusive to ALAN exposure, cellular process, localization, and pigmentation were identified as primary categories, with no circadian rhythm-related terms appearing exclusively enriched in the Midday vs. ALAN comparison (Figure 3). When comparing dorsal and ventral skin (data not shown), we observed a differential expression of OPN5, a non-visual opsin previously detected in the skin of green anoles (Trejo-Reveles, et al., submitted). Notably, OPN5 expression was differentially regulated regardless of skin exposure to artificial light at night (ALAN). Interestingly, the dorsal skin exhibited the highest levels of OPN5 expression (Appendix 3), surpassing even the expression levels found in brain tissue. There were no sex differences in gene expression patterns with most of the genes following the same expression patterns regardless of the tissue type. Key genes for each comparison are summarized in Table 1, and their respective expression patterns are found in Appendix 3. GO analysis showed significant enrichment only in the brain, where both Midday vs. ALAN and Midday vs. Midnight comparisons highlighted the regulation of circadian rhythms. No terms were significantly enriched in the skin or the liver (p-value ≤ 0.05).

**Table 1.** Key genes found differentially expressed (DE) in each tissue and the possible connection to artificial light at night (ALAN) in green anole lizards.

| Tissue | DE key genes | Possible implications |
| --- | --- | --- |
| Brain | PER1, NR1D1, NR1D2, CRY1, CRY2, RH2, FOS, GHRHR, ENSACAG00000017407 (MYL4) | Disruption of circadian regulation and potential effects on hormonal rhythms. NR1D1/NR1D2 and CRY2/CRY1 involvement in photoperiodism may affect seasonal and daily physiological processes, while RH2 expression suggests sensitivity to green light, possibly influencing testosterone levels and behavior. FOS upregulation may indicate stress or immediate early gene activation. GHRHR upregulation suggests impacts on growth hormone signaling. The novel transcripts, including MYL4, could imply emerging regulatory pathways influenced by ALAN. |
| Liver | GCG, PER1, NOCT, F-box domain-containing protein, GAD2, ACTA1, KLF9, MT-ND6 | Altered glucagon production and glucose homeostasis due to GCG and PER1 regulation, impacting metabolism and energy balance. NOCT downregulation suggests disruption in lipid metabolism and resistance to obesity, potentially leading to metabolic dysregulation. GAD2 upregulation implies involvement in neurotransmitter signaling or energy metabolism, while ACTA1 and KLF9 downregulation highlight |
|  |  | structural and transcriptional regulation disruptions under ALAN exposure. |
| Skin | GRIA2, GRM5, OPN5, PND4, NPY, SNAP25, DNER, CCDC | Increased expression of GRIA2 and GRM5 could influence circadian rhythm maintenance and sleep-wake cycles. The downregulation of OPN5 in ventral skin indicates potential roles for extra-retinal photoreceptors in light perception. PND4 involvement in wound healing and cell proliferation suggests additional responses in the skin under ALAN. The upregulation of NPY, SNAP25, and DNER may reflect roles in non-neuronal processes such as tissue remodeling, signaling pathways, or cellular interactions in the skin. CCDC downregulation highlights gene-specific suppression under ALAN |

When comparing the Night vs. ALAN conditions, we observed the highest number of differentially expressed genes (DEGs) in the skin, exceeding the numbers identified in the previous two comparisons (2023). Notably, GRIA2 was upregulated at levels similar to those observed in the ALAN vs. Midday comparison, with additional genes such as GRM5 (up regulated) also showing differential expression. New DEGs identified included NPY, SNAP25, and DNER, all of which were upregulated, while genes such as CCDC were downregulated (Appendix 4 A).

In the brain, the ALAN vs. Midnight comparison revealed only 50 DEGs, representing the lowest number among all comparisons. Notably, no genes were downregulated in this condition. Among the upregulated genes, the Growth Hormone-Releasing Hormone Receptor (GHRHR) was the annotated gene that exhibited the most striking upregulation (p-value of 7.93e-06, log fold change 2.93). Novel transcripts, including ENSACAG00000006458, ENSACAG00000011927, and ENSACAG00000017407 (this last one presumed to encode for MYL4), were also up-regulated. Other DEGs included circadian-related genes such as PER1 and interestingly FOS was also up regulated in this comparison. (Appendix 4B).

Liver tissue displayed a total of 259 DEGs, the fewest observed across the comparisons. While some genes were consistent with those identified in the Midday vs. ALAN comparison, such as CGC and MT-ND6, novel targets also emerged. Among the newly identified genes, GAD2 was upregulated, whereas ACTA1 and KLF9 were downregulated. (Appendix 4C).

### Further exploration of the dataset

This comprehensive study provides valuable insights into the genetic impacts of artificial light exposure on green anoles, highlighting significant changes in circadian rhythm and metabolic gene expression across tissues. Due to the scope and complexity of the data, it is challenging to capture and summarise all possible comparisons and interactions within a single report. To address this, we developed a publicly accessible online database to allow researchers to further explore the dataset in depth, analyse specific genes, and examine differential expression patterns tailored to additional research questions. This interactive resource is available at https://vtrejor.shinyapps.io/green_anole_rnaseq/. It provides a user-friendly platform for visualising and downloading data, supporting a more nuanced understanding of how light exposure affects genetic expression in reptiles. By making this resource available, we hope to facilitate further research, collaboration, and discovery in the field of vertebrate biology and conservation.

## Discussion

This study presents the first transcriptomics database focusing on ALAN effects in lizards, which given their distribution in urban habitats (French et al. 2018), are frequently subjected to light pollution (e.g., Thawley and Kolbe 2020; Taylor et al., 2022). This report focused on the effect of ALAN on circadian and metabolic responses in the brain and liver as well as opsin expression in the skin. To complement these findings a comprehensive publicly accessible database was developed for researchers to further explore the effects of ALAN.

In the brain, which included the pineal gland, similar patterns of DEGs under ALAN versus natural light-dark cycles were observed, which is consistent with the brain containing the master internal biological clock (Miller et al., 2015). However, some genes displayed a significant difference in log-fold changes, or were not differentially expressed at all in one of the comparisons. For example, *NR1D2* was upregulated when Midday vs. Midnight was compared; its ortholog, *NR1D1*, was downregulated in the same comparison. Under ALAN exposure, only *NR1D1* was differentially expressed. Both *NR1D1* and *NR1D2* are associated with photoperiodism, as they play a key role in the regulation of circadian rhythms by modulating gene expression in response to light. *NR1D1* (also known as REV-ERBα) and *NR1D2* (REV-ERBβ) act as transcriptional repressors within the molecular circadian clock, influencing the stability and amplitude of circadian rhythms through the repression of genes involved in metabolic, inflammatory, and behavioural pathways (Preitner et al., 2002; Bugge et al., 2012). These genes are particularly sensitive to light, which entrains their rhythmic expression patterns, providing a direct link from external light-dark cycles to internal biological rhythms (Sato et al., 2004). Although the role of these genes in the regulation of the reptile circadian rhythm is less understood, studies in other vertebrates suggest that these genes are crucial for photoperiodic responses, including those associated with feeding and seasonal reproduction (Chauvet et al., 2016). Related to light detection, one interesting finding was that *RH2*, a green light-sensitive opsin, showed differential expression in the brain when exposed to ALAN. While *RH2* is typically studied in the eye, prior data links its involvement to seasonal changes in circulating testosterone in male green-spotted grass lizard Takydromus viridipunctatus (Tseng et al., 2018. Unpublished data from our laboratory (Trejo-Reveles et al., submitted) suggest that opsins, particularly *OPN5*, vary in expression across green anole tissues. The role of opsins in response to ALAN is yet to be investigated, but this study highlights the importance of extra retinal photoreceptors in structures such as the brain, pineal gland and skin.

In the liver, *GCG*, which encodes glucagon, was differentially expressed under ALAN, suggesting that disrupted circadian cycles may alter hormone production in lizards (Martin & White, 2016). Glucagon plays a crucial role in maintaining glucose homeostasis by stimulating glycogen breakdown and glucose release, particularly during fasting states. Its secretion follows a circadian rhythm regulated by the liver’s internal clock and feedback mechanisms from other organs, such as the pancreas and the hypothalamus, aligning glucagon levels with the body’s metabolic needs over the day-night cycle (Kalsbeek et al., 2010; Vieira et al., 2015). The rhythmic release of glucagon is driven, in part, by core clock genes, including *PER1*, and is influenced by light exposure, which can disrupt glucagon’s circadian oscillations, leading to altered glucose metabolism and potential metabolic imbalance (Grunst et al., 2023). This regulation is essential for energy balance, as glucagon levels peak during nocturnal phases, preparing the body for fasting periods, and are suppressed during feeding periods (Guan & Lazar, 2022). *PER1* was differentially expressed under both conditions, indicating that altered light-dark cycles could affect clock gene expression in the liver, potentially as a result of an imbalanced glucagon regulation (Ando et al., 2013). In addition, *NOCT*, which is involved in the liver circadian clock and lipid metabolism (Kulshrestha et al., 2023), was downregulated under natural light-dark cycles. Knockout studies in mice have demonstrated *NOCT*’s roles in cellular differentiation and metabolism, with implications for higher energy expenditure, and reduced adiposity (Le et al., 2019; Abshire, et al., 2020).

In the skin, many genes were upregulated under ALAN exposure. Notable genes include *GRIA2*, which is associated with circadian rhythm maintenance in the mouse brain, and *GRM5*, which is involved in sleep-wake cycles (Raap et al., 2015; Taylor et al., 2022). Genes that were differentially expressed in normal light-dark cycles in the skin were generally not associated with circadian rhythms, except for *NR1D1*. Instead, genes such as *PNK4*, involved in wound healing, and *CEBPD*, related to cell death and cell proliferation (Balamurugan & Sterneck 2013) were differentially expressed. Our previous unpublished data (Trejo-Reveles et al., submitted) have shown that *OPN5* expression is downregulated in ventral skin exposed to light compared to dorsal skin. These data presented in this study confirm that *OPN5* is down-regulated in ventral skin regardless of ALAN exposure. This suggests a putative role of extra retinal photoreceptors in the green anole and highlights the importance of natural light-dark cycles.

Several reptilian studies have focused on the effects of artificial light exposure on behaviour, but have not specifically identified genetic markers to indicate stress or triggers for those behavioural changes. Lizards, which are common in the wild including urban environments, face significant challenges from light pollution. For example, previous studies using a diversity of lizard species have shown that nocturnal activity and foraging levels are increased under ALAN, but this is generally associated with reduced performance during the day (e.g., Martin et al. 2018, Mauer et al. 2019, Oda et al. 2020, Kolbe et al. 2021). Another study showed that green anoles exposed to ALAN were more active at night, using the nocturnal artificial light to explore, forage, and display. During the day, ALAN exposed lizards exhibited reduced activity, and displayed increased fat pad and testes sizes, suggesting shifts in metabolic and reproductive processes (Taylor, et al., 2022). Our findings align with these observations; for instance, genes such as *GRM5*, associated with nocturnal activity, were upregulated under ALAN exposure. ALAN has been shown to have no effect on green anole offspring quality (Clark et al., 2017), however clear ALAN induced changes in adult lizard ovaries was observed as *TTR* was upregulated in this study (data not shown). *TTR* accumulates in the choroid plexus during the dark phase of the circadian rhythm, and it is known to be influenced by sex, age and circadian rhythms. In mice, *TTR* plays a role in preparing the uterus for embryo implantation potentially influencing offspring quality (Fame et al., 2023; Duarte et al., 2020; Diao et al., 2010).

In conclusion, this study provides the first transcriptomic analysis to examine the effects of ALAN on a reptile species. In addition, these data provide a valuable public resource for researchers interested in the effects of light pollution on reptiles. The findings reveal that ALAN exposure disrupts the expression of key circadian and metabolic genes across tissues, highlighting the sensitivity of these lizards to light pollution. Specifically, the differential expression of clock-related genes such as *PER1*, *NR1D1*, and *CRY2*, alongside glucagon-related genes in the liver, underscores the influence of artificial light on fundamental physiological processes, from circadian rhythm regulation to glucose homeostasis. The observed modulation of photoreceptor genes, particularly in skin and brain, provides evidence of extra-retinal photoreception in lizards and suggests an adaptive response that may be crucial for these reptiles to cope with light-polluted environments. While these findings provide a comprehensive understanding of how ALAN exposure affects the gene expression profile in green anoles, they also emphasise the need to maintain natural light-dark cycles in wild urban habitats to support optimal physiological functioning. Investigating the functional roles of genes such as *OPN5* and *RH2* in extra-retinal and retinal photoreception, and their potential contribution to behavioural and physiological adaptations in light-polluted environments, will deepen our understanding of photoreception beyond ocular tissues. Furthermore, comparative studies across reptile species exposed to ALAN could reveal evolutionary adaptations to light pollution, offering insights into the resilience of reptile populations in urbanised landscapes. Longitudinal studies tracking gene expression changes across life stages will elucidate whether prolonged exposure to ALAN induces cumulative genetic or phenotypic changes, particularly concerning reproductive fitness and stress responses. Future studies will embrace a transcriptomic approach to further investigate the broader impacts of ALAN on reptilian biology. Indeed, ATACseq, and single cell sequencing, will be key to the understanding which cell populations are important for physiological and behavioural adaptation in our rapidly illuminated world. These findings will be instrumental for conservation efforts aimed at mitigating light pollution and preserving natural light cycles, which are integral to the health and survival of animals in their natural habitats.

## Acknowledgements

This work was supported by an International Institutional Award to the University of Edinburgh (BB/Y51410X/1) and Roslin Institute Strategic Grant (BBS/E/RL/230001C) funding from the UK Biotechnology and Biological Sciences Research Council to Simone L. Meddle along with financial support from the Trinity University Office of Academic Affairs to Michele A. Johnson. We would like to thank Dale Cochran and members of the Johnson Lab for all of their fantastic help in the laboratory and field. Green anole collection was performed under Scientific Permit SPR-0310-045 to Michele A. Johnson from Texas Parks and Wildlife Department, with approval from Trinity University’s Animal Research Committee, protocol 051122-MJ2.

## Author contributions

V.T-R, M.A.J, and S.L.M. designed the research; V.T-R, M.A.J, G.E.A, F.J.P and A.R.J. performed the research and V.T-R, M.A.J, and A.R.J. analysed the data; V.T-R, M.A.J, and S.L.M. wrote the manuscript. All the authors were involved in drafting and revising the manuscript.

## Abbreviation list

ALAN: Artificial light at night
DEG: Differentially expressed genes GO - Gene ontology
UVB: Ultraviolet B (light) RNA - Ribonucleic acid
RIN: RNA integrity number FASTQC - Fast quality control
PCA: Principal component analysis

## Appendix Figure Legends

**Appendix 1.**
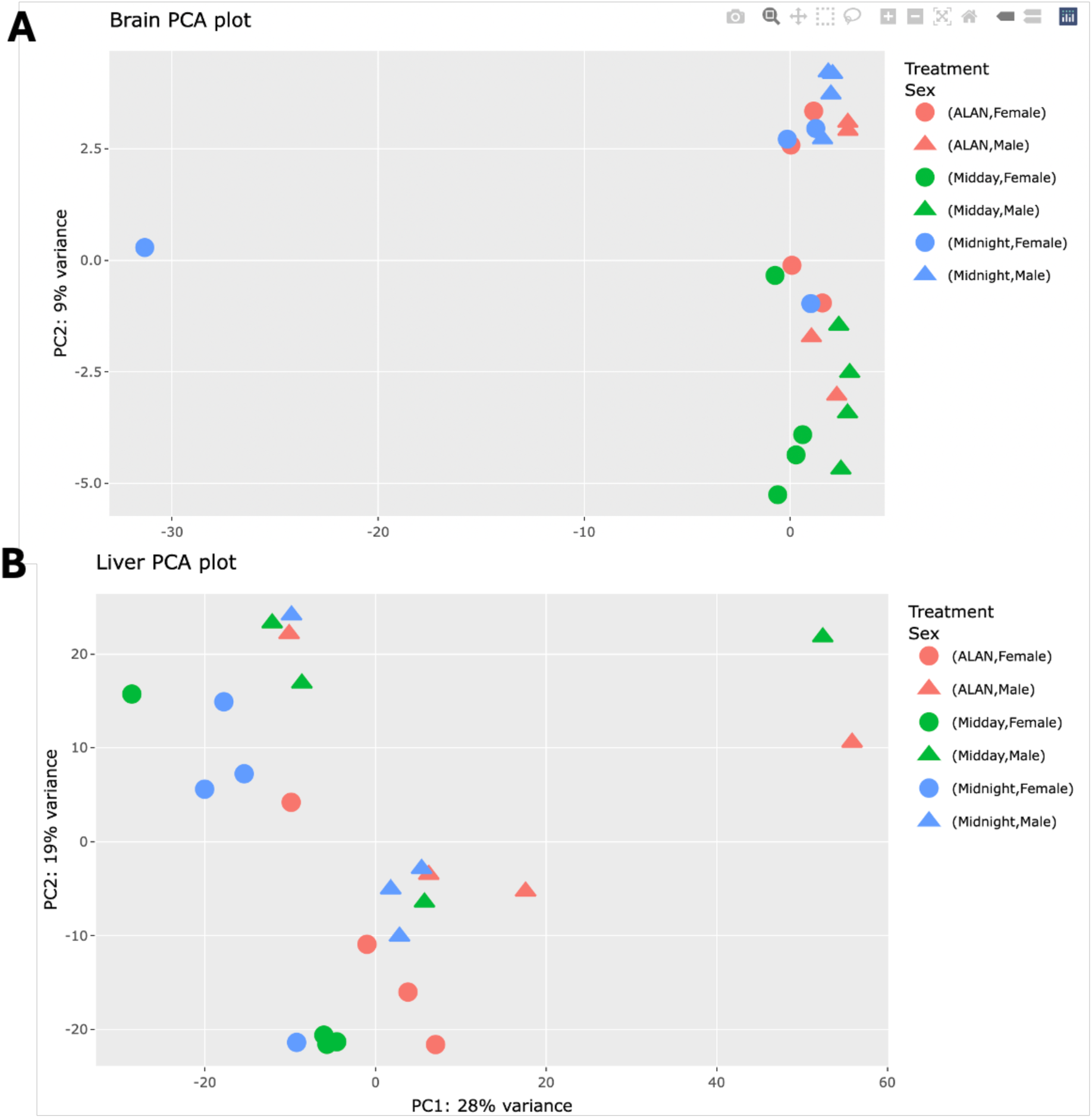
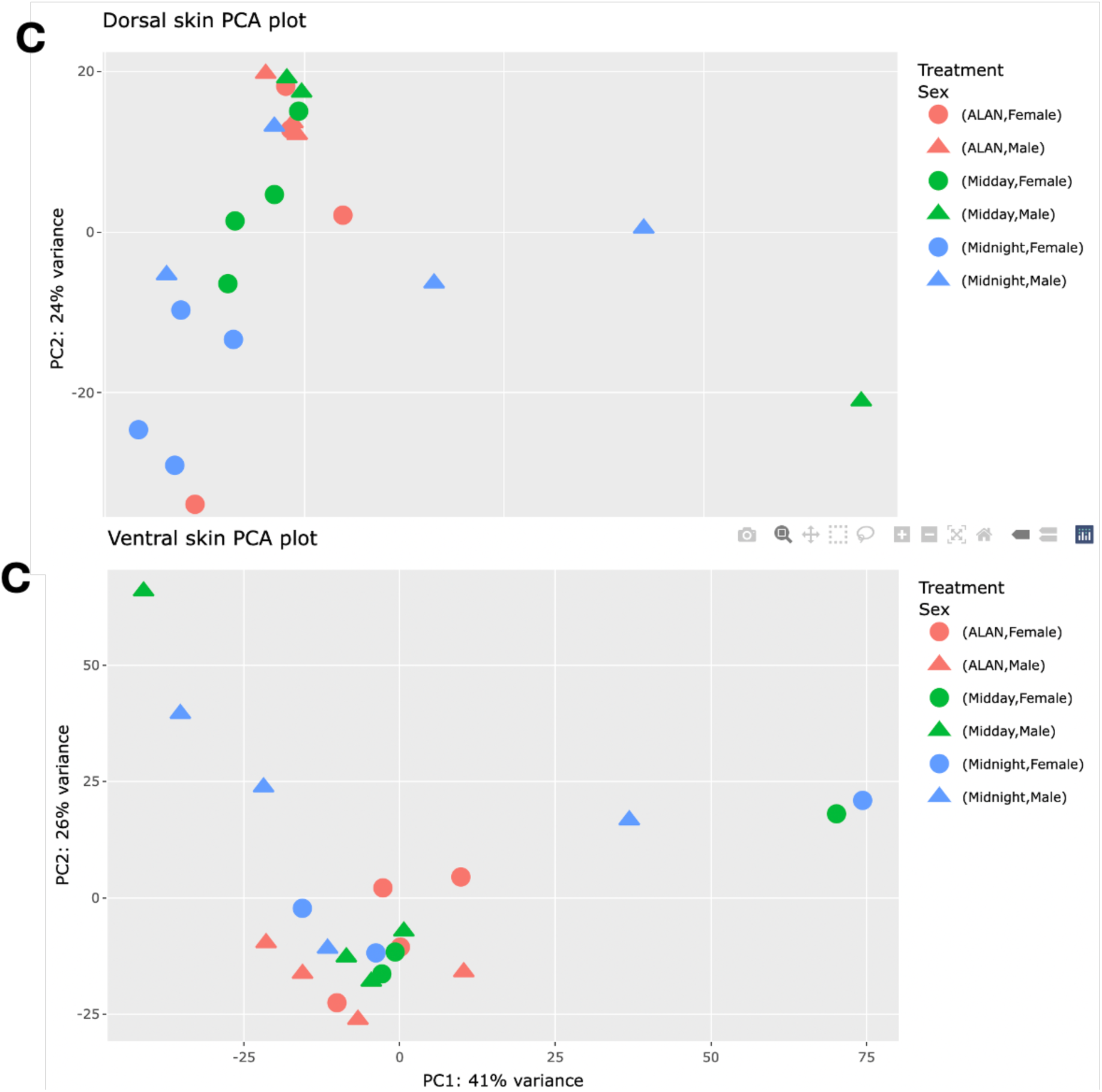
PCA plots of individual tissues from green anoles; the colour depicts the treatment and the shape depicts the sex. None of the PCA plots show clear clustering between different lighting conditions or sex.

**Appendix 2.**
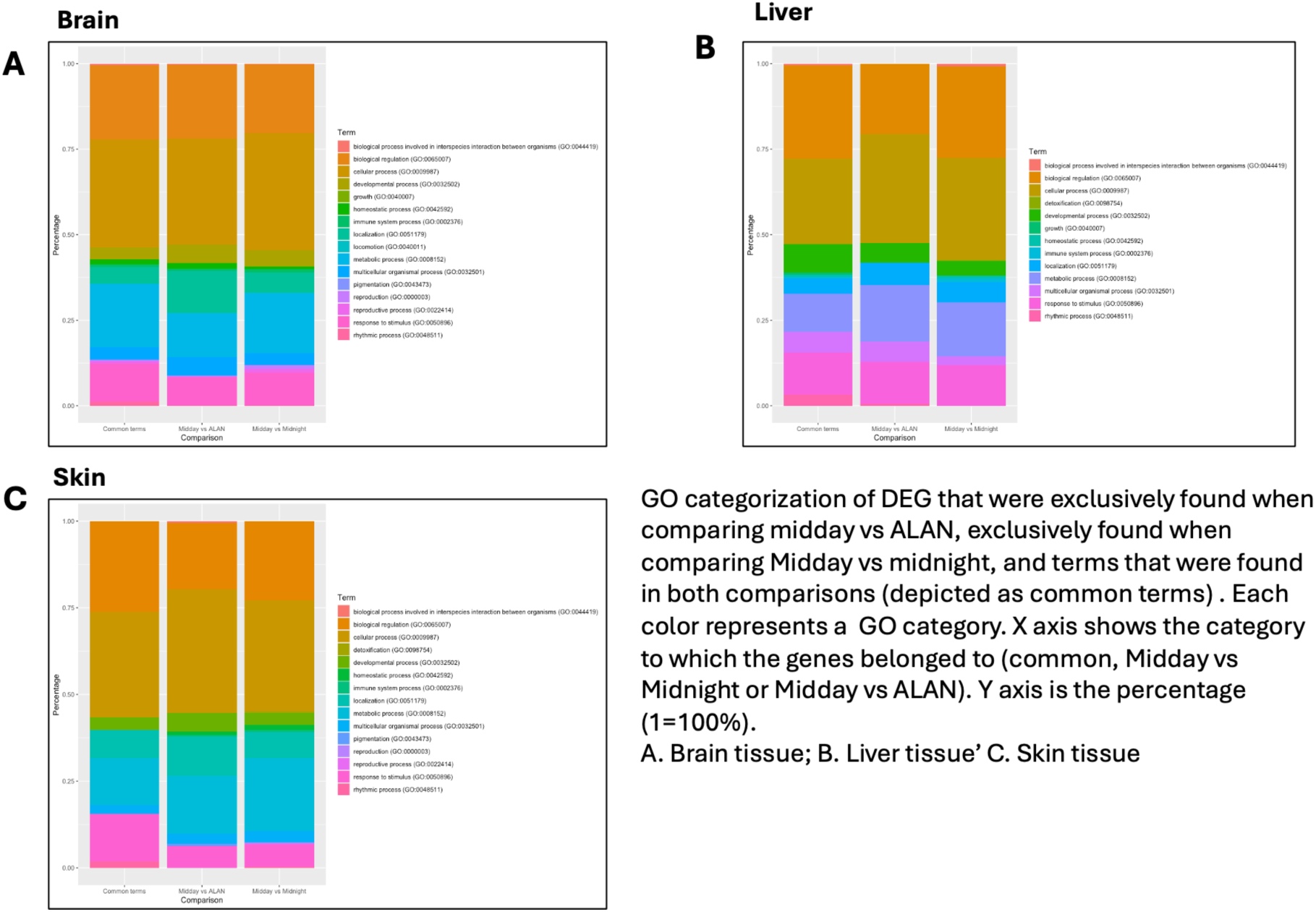
GO categorization of DEG that were exclusively found when comparing Midday vs ALAN, exclusively found when comparing Midday vs Midnight, and terms that were found in both comparisons (depicted as common terms). Each colour represents a GO category. X axis is the category to which the genes belonged to (common, Midday vs Midnight or Midday vs ALAN). Y axis is the percentage (1=100%). **A**. Brain; **B**. Liver **C**. Skin.

**Appendix 3.**
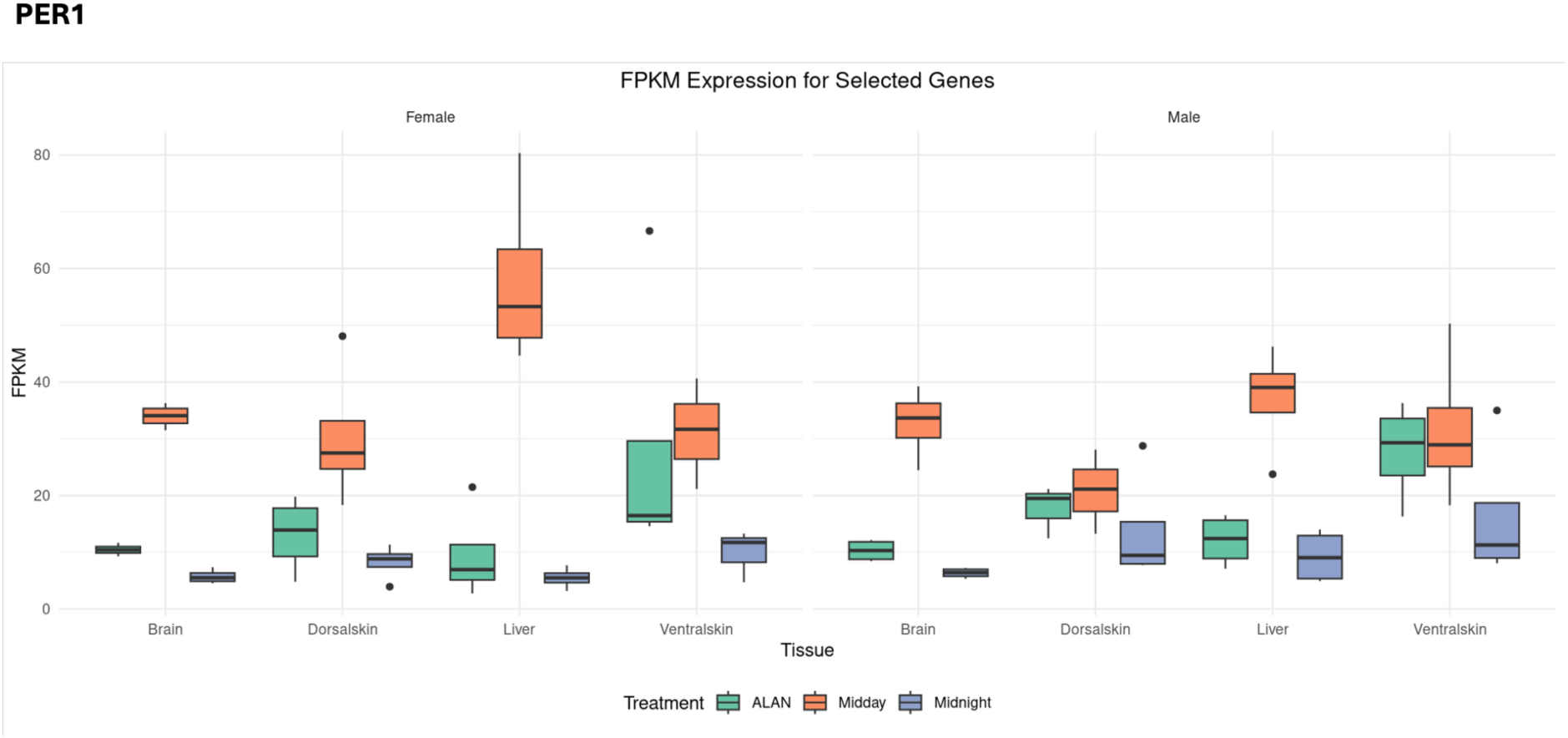
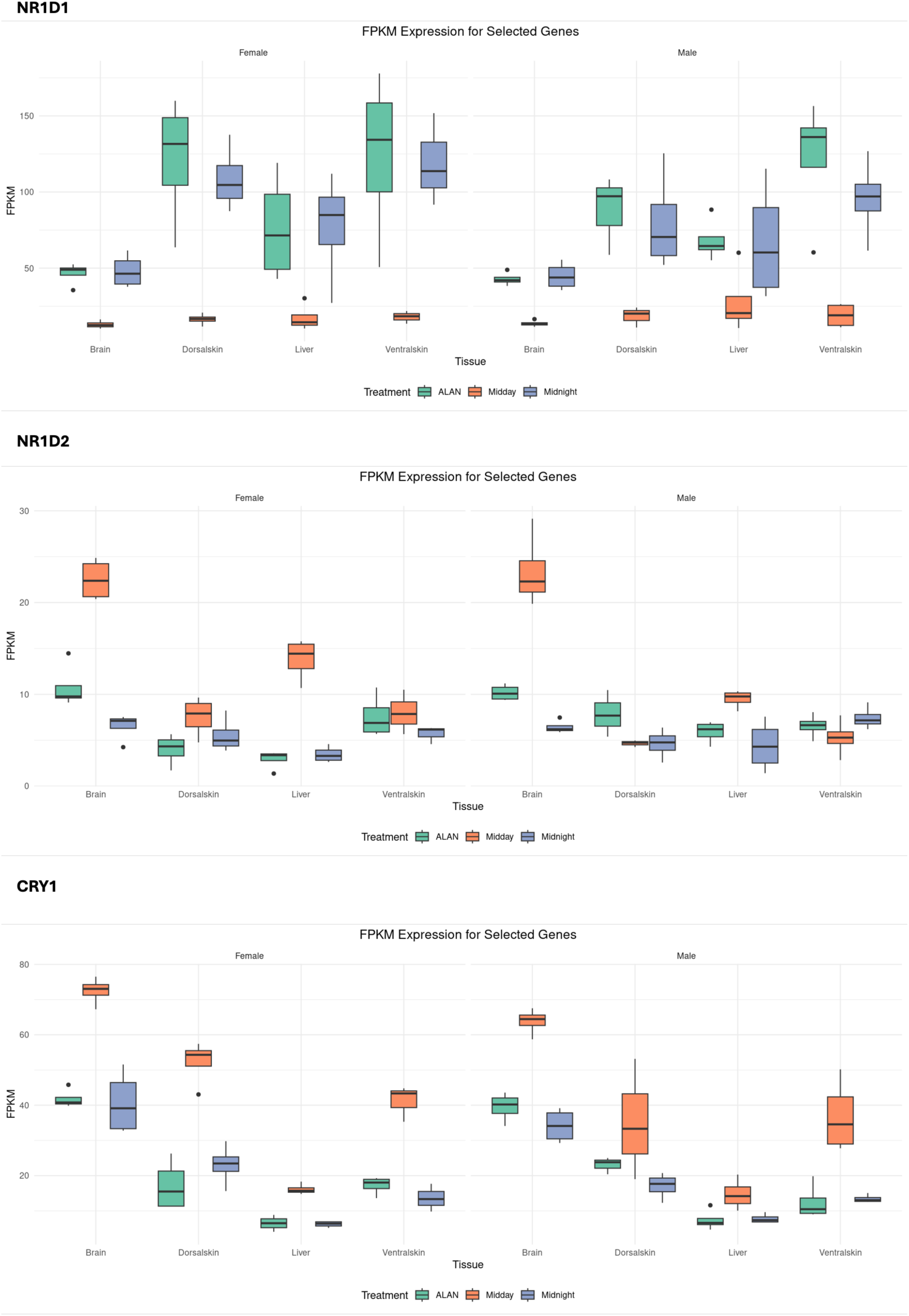
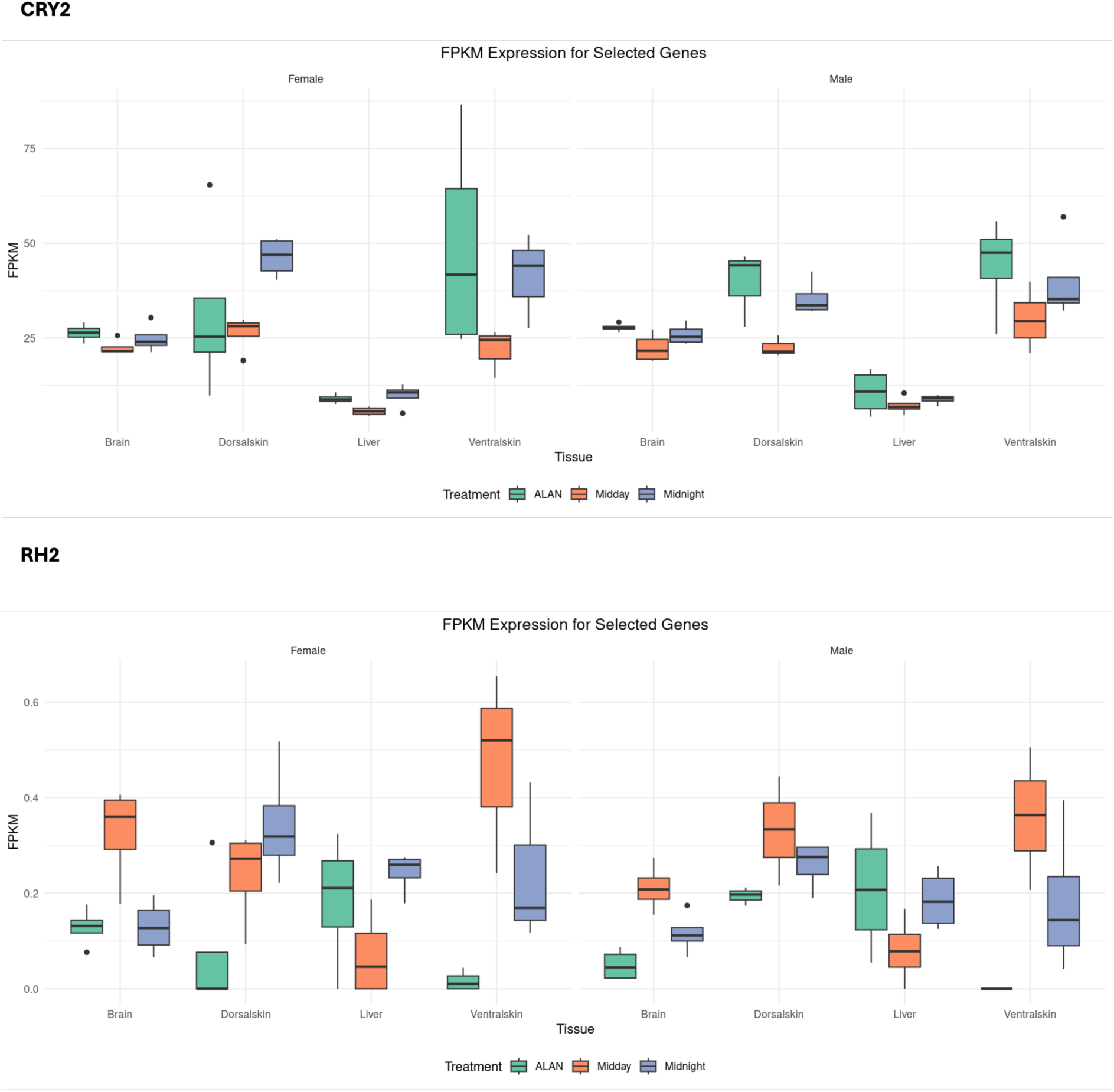
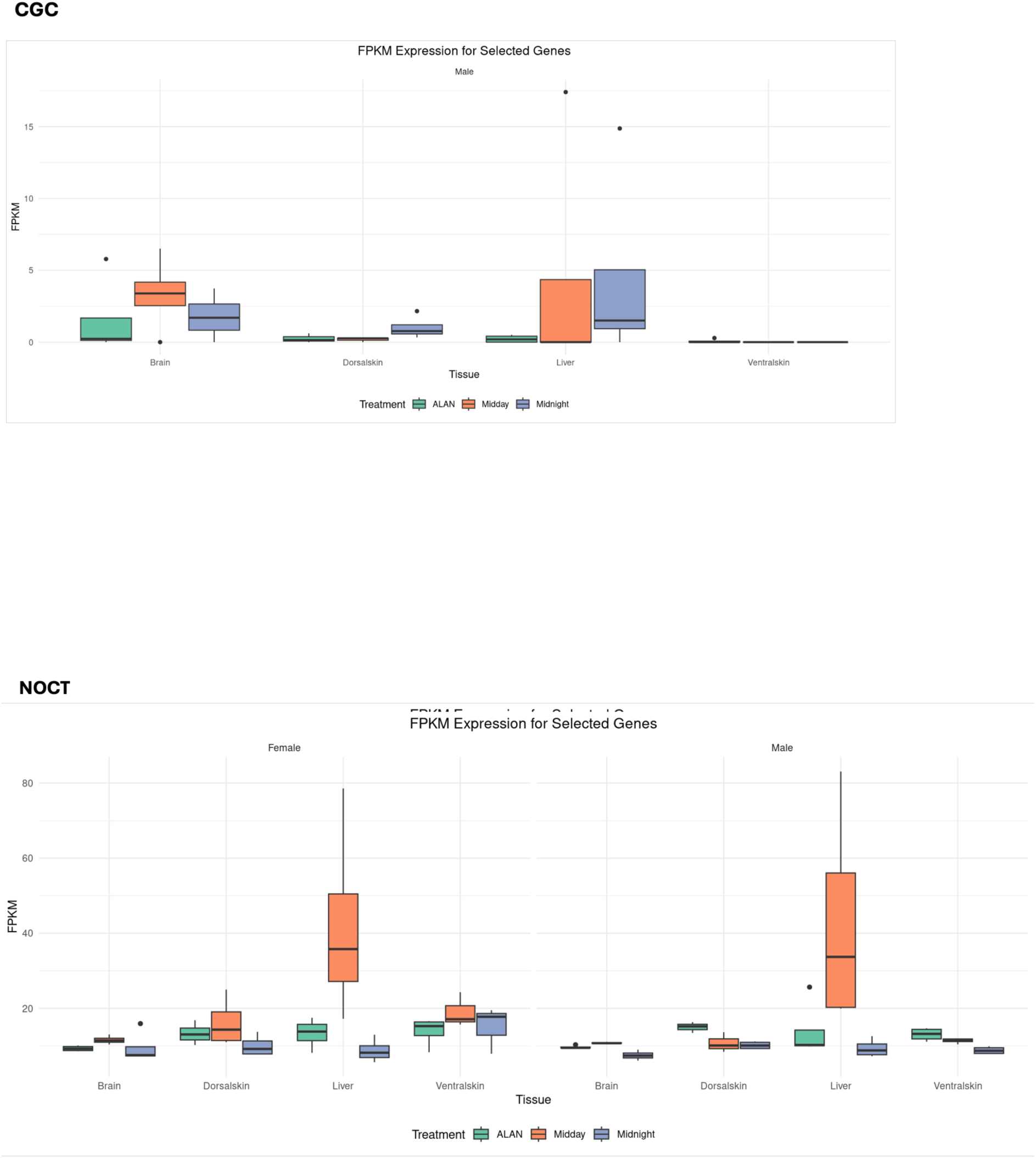
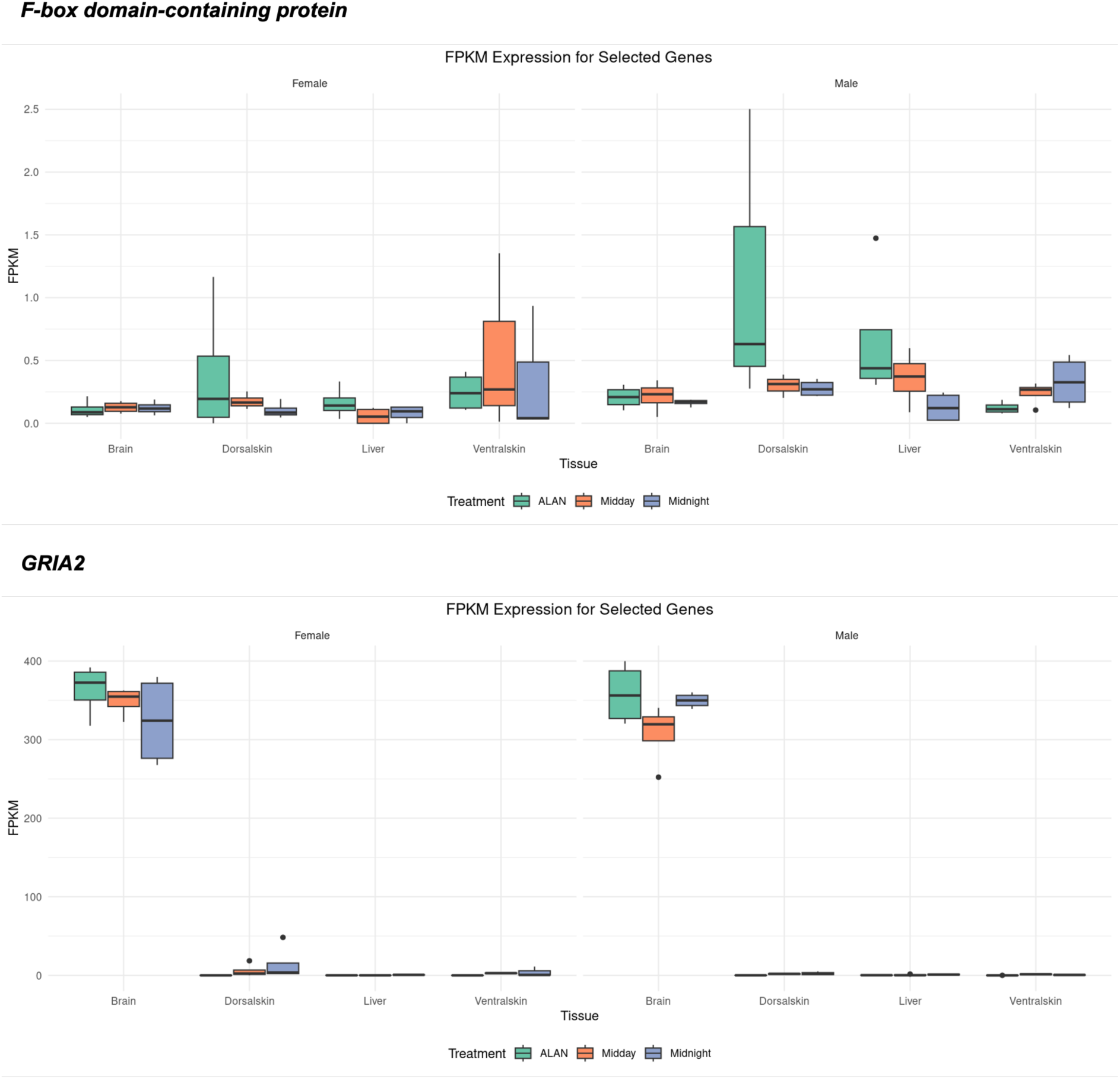
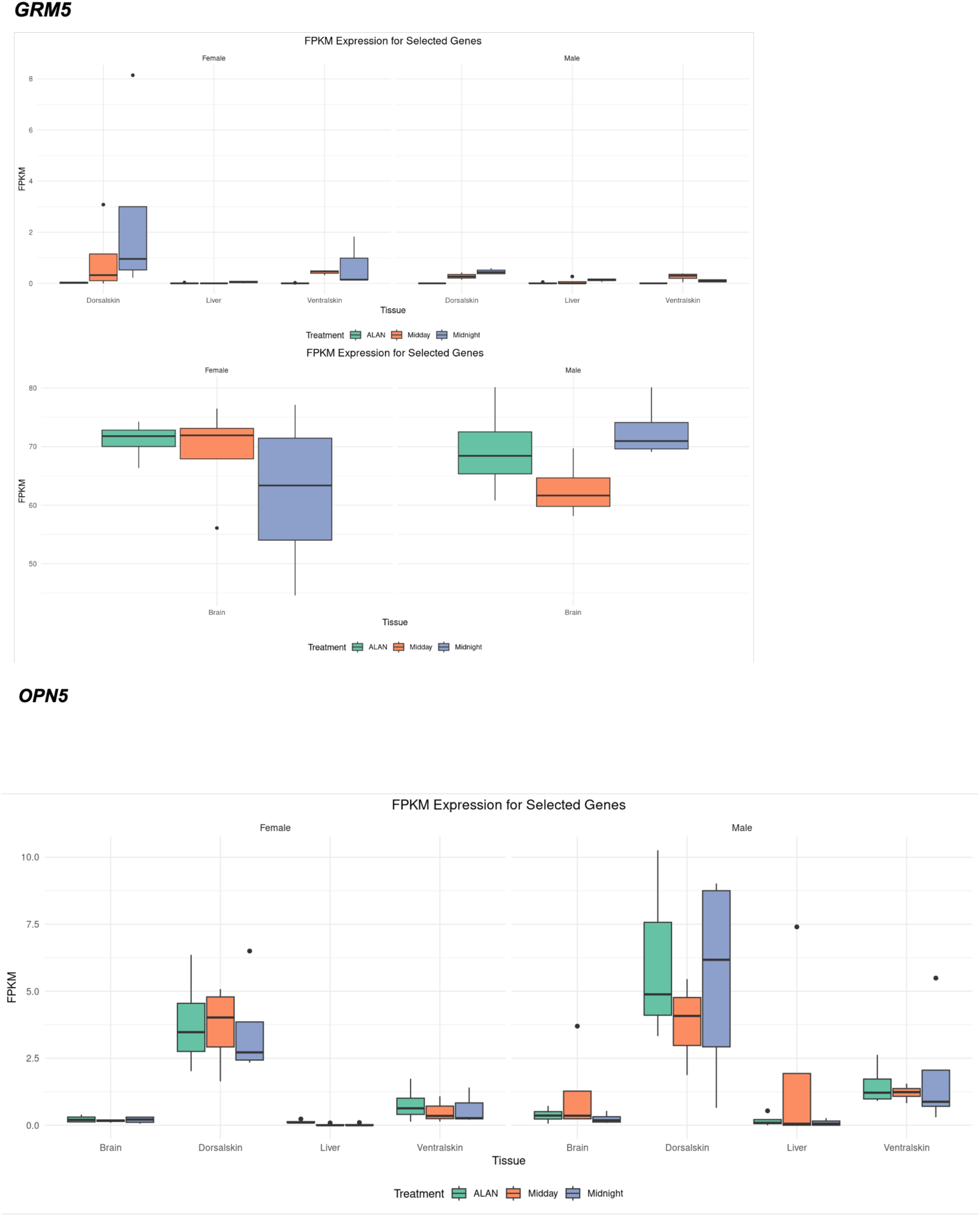
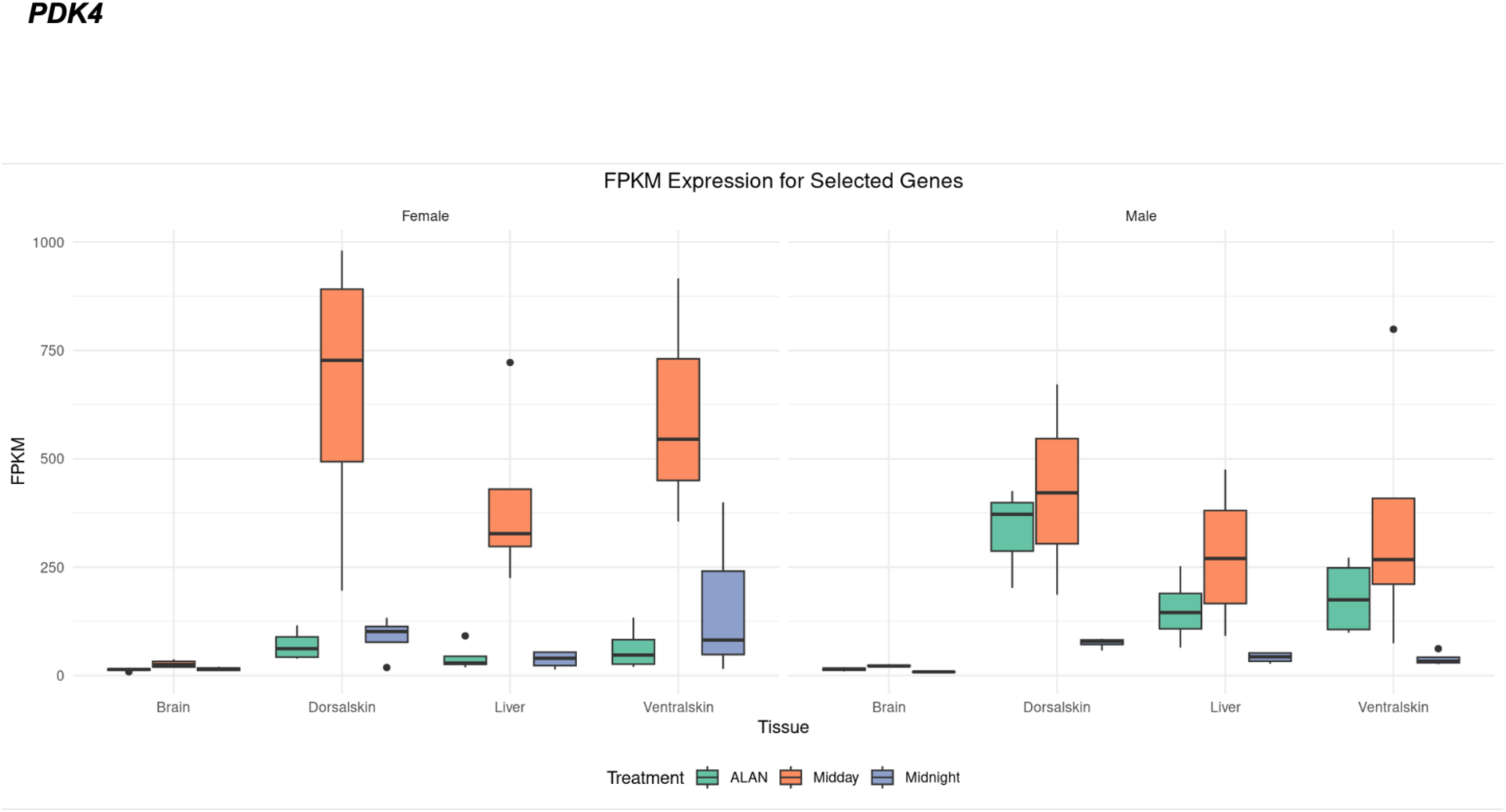
FPKM values of genes of interest in different tissues, conditions and sex. Each boxplot represents a light exposure condition. X axis represents the tissue; Y axis depicts the FPKM and colours represent the lighting condition. Female and male individuals’ values are shown next to each other.

**Appendix 4.**
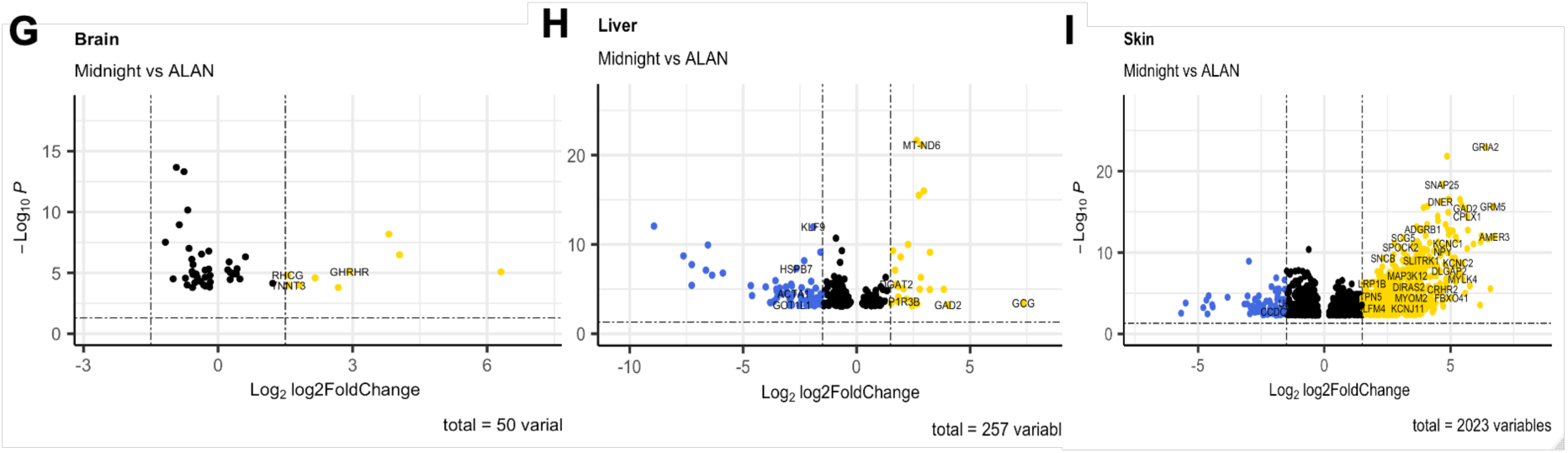
ALAN vs Midnight comparison volcano plots

